# CO_2_ per se activates carbon dioxide receptors

**DOI:** 10.1101/787481

**Authors:** Pingxi Xu, Xiaolan Wen, Walter S. Leal

## Abstract

Carbon dioxide has been used in traps for more than six decades to monitor mosquito populations and help make informed vector management decisions. CO_2_ is sensed by gustatory receptors (GRs) housed in neurons in the maxillary palps. CO_2_-sensitive GRs have been identified from the vinegar fly and mosquitoes, but it remains to be resolved whether these receptors respond to CO_2_ or bicarbonate. As opposed to the vinegar fly, mosquitoes have three GR subunits, but it is assumed that subunits GR1 and GR3 form functional receptors. In our attempt to identify the chemical species that bind these receptors, we discovered that GR2 and GR3 are essential for receptor function and that GR1 appears to function as a modulator. While *Xenopus* oocytes coexpressing *Culex quinquefasciatus* subunits CquiGR1/3 and CquiGR1/2 were not activated, CquiGR2/3 gave robust responses to sodium bicarbonate. Interestingly, CquiGR1/2/3- coexpressing oocytes gave significantly lower responses. That the tertiary combination is markedly less sensitive than the GR2/GR3 combination was also observed with orthologs from the yellow fever and the malaria mosquito. By comparing responses of CquiGR2/CquiGR3- coexpressing oocytes to sodium bicarbonate samples (with or without acidification) and measuring the concentration of aqueous CO_2_, we showed that there is a direct correlation between dissolved CO_2_ and receptor response. We then concluded that subunits GR2 and GR3 are essential for these carbon dioxide-sensitive receptors and that they are activated by CO_2_ per se, not bicarbonate.

## INTRODUCTION

Carbon dioxide is the oldest known and the most powerful mosquito attractant [1]. It has been widely used for decades for trapping blood-seeking female mosquitoes [2, 3] to monitoring populations and to assist in integrated vector management programs. In addition to being an attractant sensu stricto, CO_2_ gates mosquito perception of other sensory modalities [4–6]. In the vinegar fly, *Drosophila melanogaster,* CO_2_ elicits through different pathways attraction and aversion behaviors [7]. Mosquitoes sense CO_2_ with olfactory receptor neurons (ORNs) [8–10] housed in peg sensilla (= capitate pegs) in the maxillary palps [11]. It has been unambiguously demonstrated that CO_2_ is sensed by the gustatory receptors DmelGR21a and DmelGR63a, housed in antennal neuron ab1C of the vinegar fly [12–14], but it remains to be resolved whether carbon dioxide receptors are activated by CO_2_ per se or bicarbonate [13–16].

Mosquitoes have three closely related homologs of DmelGR21a and DmelGR63a [17], GR1, GR2, and GR3. Because they were identified before the new nomenclature for these GR genes was proposed [17], GR1-3 in the malaria mosquito, *Anopheles gambiae*, are still named AgamGR22, AgamGR23, and AgamGR24, respectively [18]. While it has been clearly demonstrated that GR3 is essential for CO_2_ reception in the yellow fever mosquito, *Aedes aegypti* [5], it is not yet known whether GR1 and GR2 are functionally redundant [5]. While no response to CO_2_ was recorded when both AgamGR22 and AgamGR24 (one copy of each) were coexpressed in the empty neuron system of the vinegar fly [19], by increasing the dosage of both transgenes by two-fold led to significant response [20]. Additionally, flies carrying one copy of each of the three subunits showed a significant response to CO_2_, albeit not as strong as responses recorded from the flying carrying two-fold of AgamGR22 and 24 [20]. Thus, AgamGR23 was implicated in CO_2_ reception in the malaria mosquito, but its role was not clarified. On the other hand, by using RNAi combined with behavioral measurements, it has been suggested that AaegGR2 had no role in CO_2_ reception by the yellow fever mosquito [21]. In summary, it remains to be determined whether all three GR subunits are functionally required for CO_2_ detection in mosquitoes or whether GR1 and GR2 are functionally redundant [5].

We envisioned that having a functional carbon dioxide-detecting system in an aqueous environment as in the *Xenopus* oocyte recording system would allow us to address with the Le Chatelier’s principle the long-standing question whether carbon dioxide receptors are activated by CO_2_ per se or by bicarbonate [13–16]. Because dissolved CO_2_ forms an equilibrium with bicarbonate in water 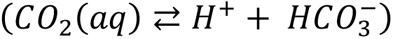, by shifting the equilibrium towards dissolved CO_2_ would increase a receptor response if CO_2_ per se binds carbon dioxide receptors, whereas a reduced response would indicate activation by bicarbonate. Here, we show that CO_2_ per se, not bicarbonate, activates the receptors. While preparing to address this question, we discovered that the gustatory receptor CquiGR2 from the southern house mosquito, *Culex quinquefasciatus*, is essential for function, whereas CquiGR1 appears to be a modulator. Additionally, we show a similar GR1 effect in the *Ae. aegypti* and *An. gambiae* CO_2_-sensing system. Specifically, CO_2_ elicited weaker responses in the tertiary receptor systems than in the respective binary counterparts devoid of GR1 (GR22 in the case of the malaria mosquito receptors).

## RESULTS AND DISCUSSION

### *Cx. quinquefasciatus* Gustatory Receptor GR2/GR3 is Sensitive to Carbon Dioxide

We cloned three gustatory receptors from the southern house mosquito, specifically CquiGR1 (VectorBase, CPIJ006622), CquiGR2 (GenBak, MN428502), and CquiGR3 (MN428503), which are likely to be involved in CO_2_ reception. Initially, we coexpressed combinations of these three GRs in the *Xenopus laevis* oocytes and recorded their responses to sodium bicarbonate. Of note, the application of sodium bicarbonate increases the sodium concentration in the perfusion Ringer, thus causing inward currents through endogenous oocyte channels due to an increase in the extracellular concentration of Na^+^ (Figure 1A, Figure S1). We, therefore, applied equivalent doses of NaCl and NaHCO_3_ in all experiments to control for the Na^+^ response. Equal responses to NaCl and NaHCO_3_ at the same dose, as in the case of naked oocytes (Figure 1A), indicate that the currents were generated by Na^+^ ions. Surprisingly, oocytes coexpressing CquiGR1 and CquiGR3 did not respond to bicarbonate as the response to sodium bicarbonate did not differ from the response to sodium chloride (Figure 1B, Figure S1). A lack of specific response to sodium bicarbonate was also observed with CquiGR1/CquiGR2- coexpressing oocytes (Figure 1C, Figure S1). By contrast, CquiGR2/CquiGR3-expressing oocytes gave robust dose-dependent responses to sodium bicarbonate (Figures 1D, 2, Figure S1). Although it is not surprising that injections of sodium chloride generated dose-dependent currents, they were significantly smaller (t-test, *P*=0.0008) than those generated by sodium bicarbonate at the same dose (Figure 2).

**Figure 1.**
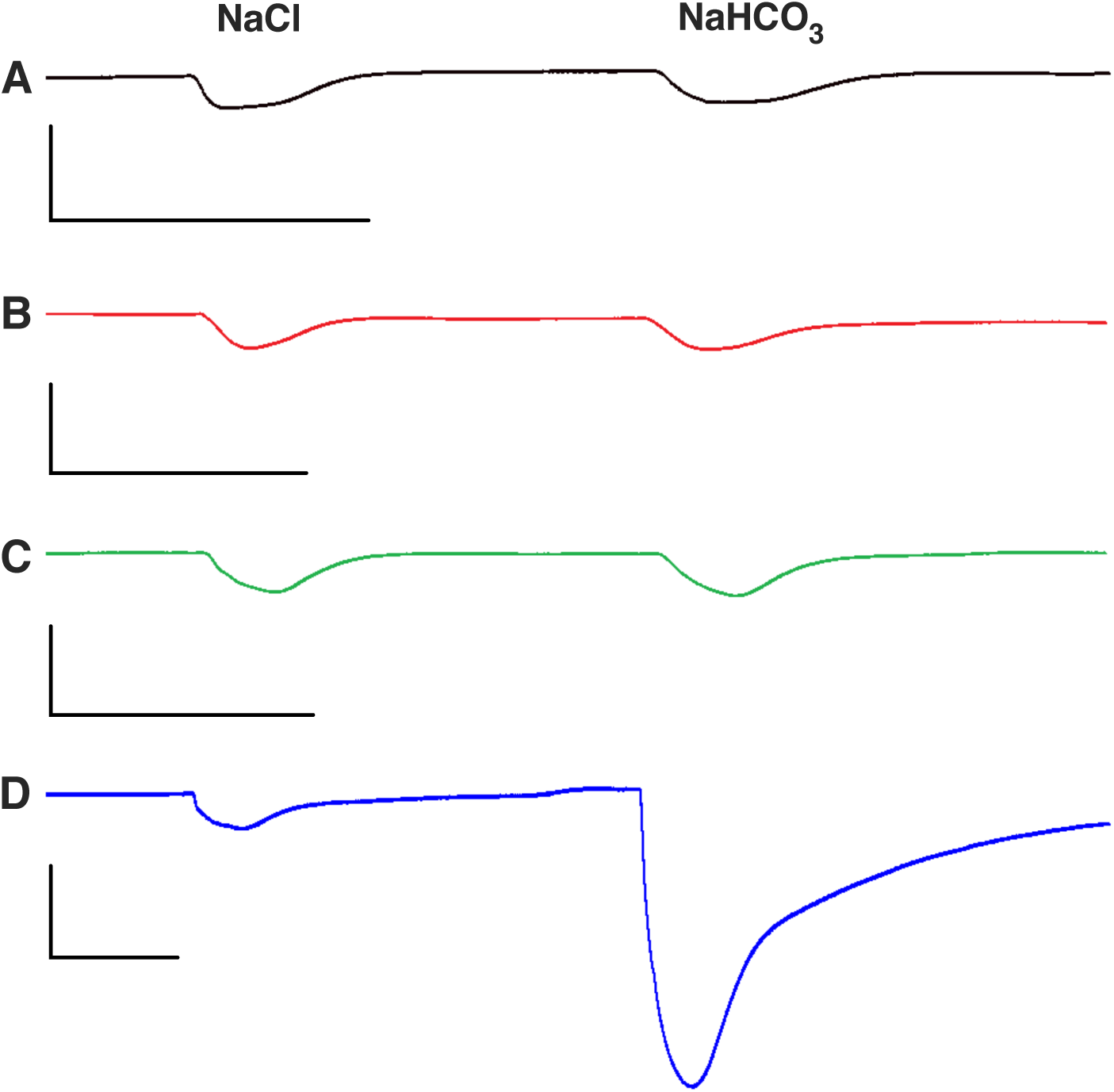
Responses of Oocytes to Sodium Chloride and Sodium Bicarbonate. Traces obtained with the following types of oocytes from the same batch and age: (A) Intact (naked) *Xenopus* oocytes. (B) Oocyte coexpressing CquiGR1 and CquiGR3 (C) Oocyte coexpressing CquiGR1 and CquiGR2 (D) Oocyte coexpressing CquiGR2 and CquiGR3. All scales are 200 nA and 0.5 min. Oocytes were first challenged with 200 mM NaCl and then 200 mM NaHCO_3_.

**Figure 2.**
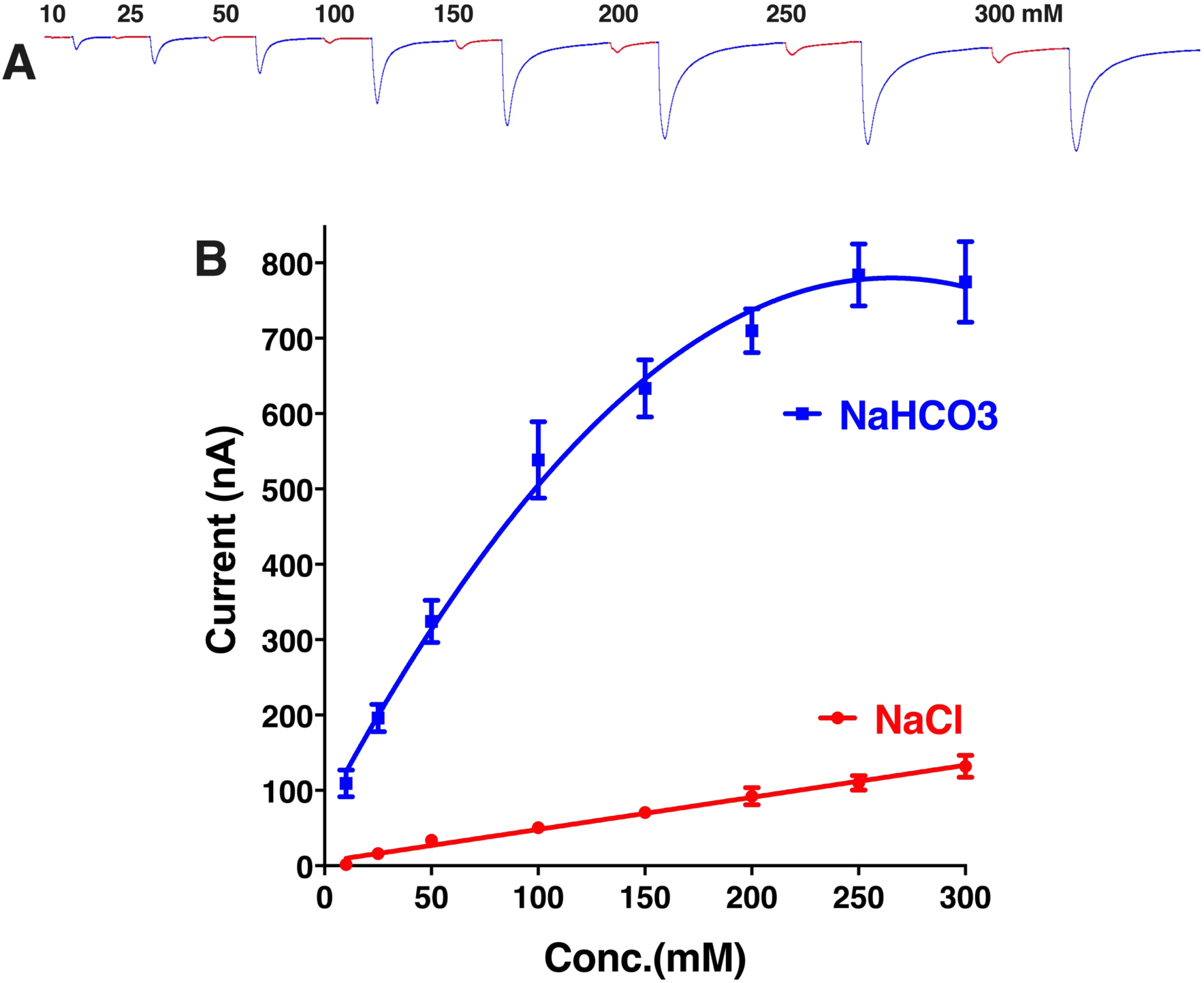
Responses of CquiGR2+CquiGR3-coexpressing Oocytes to Sodium Chloride and Sodium Bicarbonate. (A) Trace from a single oocyte preparation stimulated with increasing doses (10 – 300 mM) of the two compounds. For clarity, responses to sodium chloride and sodium bicarbonate were colored red and blue, respectively. (B) Concentration-dependent curves obtained with five different oocytes coexpressing CquiGR2 and CquiGR3. Data are presented as mean ± SEM. Some error bars for NaCl curve do not appear, because they are shorter than the size of the symbol. From left to right: 0.81, 2.59, 3.24, 5.35, 6.92, 11.24, 9.59, and 14.42.

We then compared CquiGR2/CquiGR3 and CquiGR1/CquiGR2/CquiGR3 (Figure 3). The tertiary receptor system showed a remarkably lower (t-test, *P*=0.0320) response to sodium bicarbonate than the binary receptor (Figure 3).

**Figure 3.**
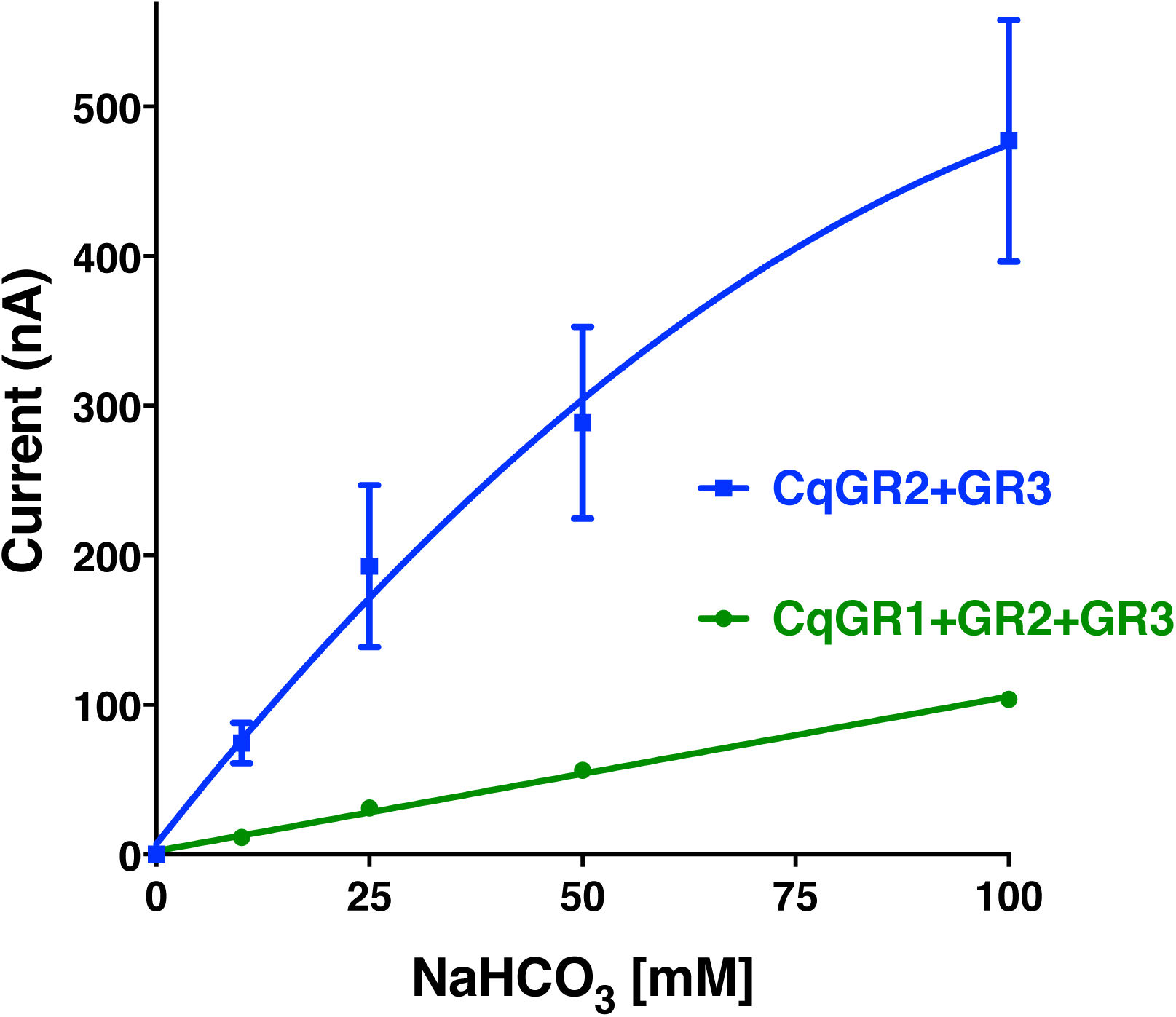
Comparative Concentration-dependent Curves for Oocytes Coexpressing either two or three Functional Subunits. Results were obtained with four oocytes coexpressing CquiGR2 and CquiGR3 and four oocytes coexpressing the three GRs. For simplicity, in figures, we use two-letter symbols for proteins as opposed to four-letter symbols (eg, Cq = Cqui) used elsewhere. Data are presented as mean ± SEM. Error bars for the green curve do not appear because they are shorter than the size of the symbol. From left to right: 0, 0.63, 3.24, 1.65, and 3.84.

Next, we examined whether variants of CquiGR1 would have different effects. We cloned five CquiGR1 variants, one of them showed an amino acid sequence identical to the sequence appearing in VectorBase (CPIJ 006622). We named this variant the wild type form. Three other variants differed in one amino acid residue each, specifically CquiGR1K456N (GenBank, MN418391), CquiGR1M365V (MN418394), and CquiGR1L246F (MN418393). Lastly, one variant differed in two amino acid residues, ie, CquiGR1D143G;R434K (MN418392). We then expressed variants along with CquiGR2+CquiGR3 in the *Xenopus* oocyte recording system and compared their responses to those obtained with CquiGR2/CquiGR3-coexpressing oocytes when stimulated with the same dose of sodium bicarbonate (Figure S2). None of the five tertiary receptor systems showed a stronger response than those recorded from CquiGR2/CquiGR3-coexpressing oocytes (Figure 4). Responses recorded from CquiGR1D143G;R434K were not significantly different from those measured from the binary receptor system (n *=* 4, unpaired t-test, *P*=0.0710). All other variants, including the wild type, generated significantly lower responses than those recorded from CquiGR2/CquiGR3- coexpressing oocytes (Figure 4). These findings suggest that CquiGR1 may modulate the receptor system response to sodium bicarbonate.

**Figure 4.**
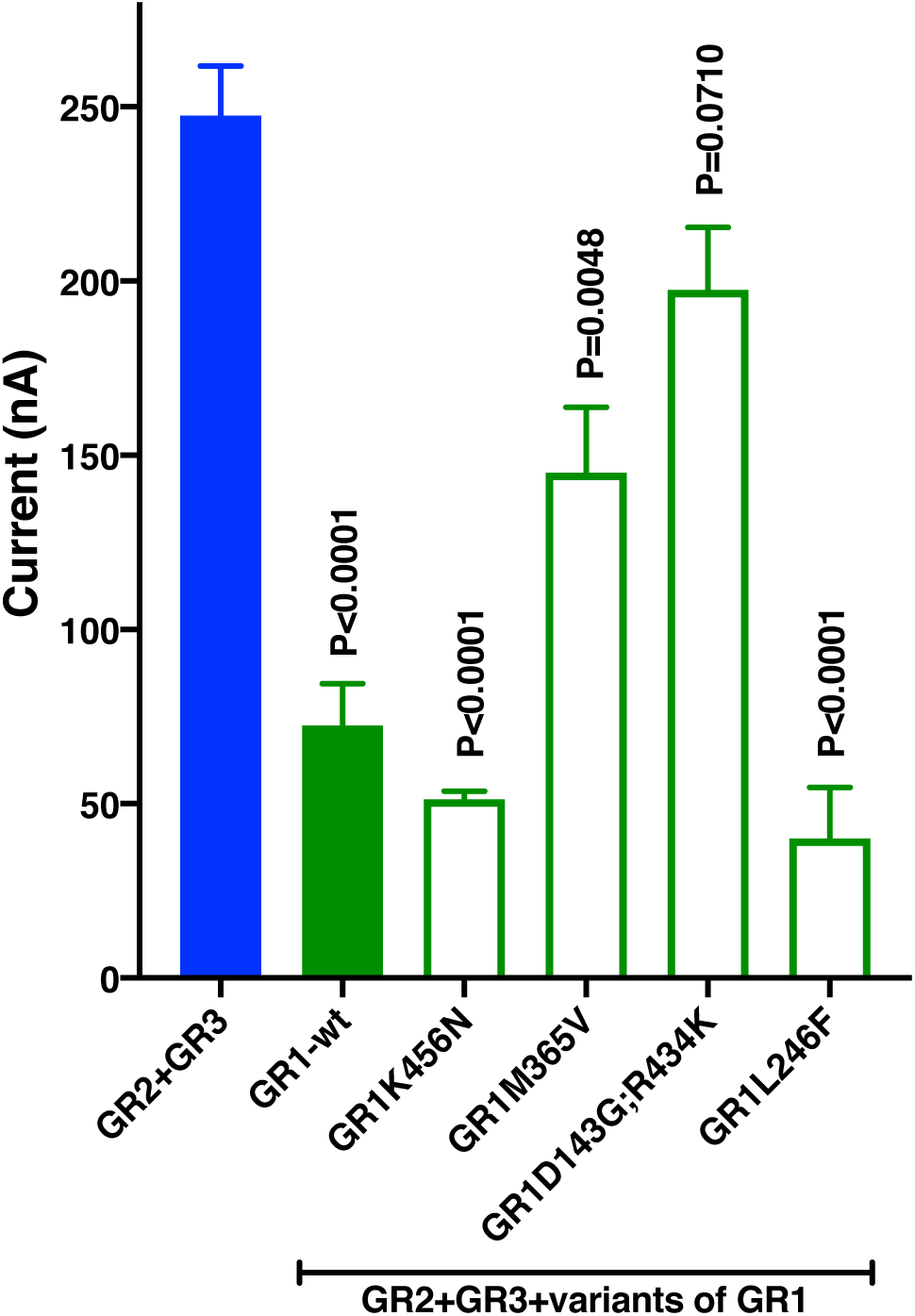
Effect of Different Variants of CquiGR1 on the Response of CquiGR2+CquiGR3- coexpressing Oocytes. All experiments were performed with four replicates with oocytes from the same batch coexpressing either CquiGR2+CquiGR3 or CquiGR2+CquiGR3 plus one of the CquiGR1 variants (K456N, M365V, D143G;R434K, and L246F). The clone with sequence identical to that in VectorBase (CJPI 006622) was named wild type (wt). Responses recorded from CquiGR2+CquiGR3-coexpressing oocytes and those recorded from oocytes coexpressing CquiGR2+CquiGR3 plus one CquiGR1variant after stimulus with 200 mM sodium bicarbonate (n *=* 4) were compared by unpaired t-test. P values are presented on the top of each variant column. Data are presented as mean ± SEM.

### Gustatory Receptors are Activated by Carbon Dioxide per se, not Bicarbonate

To address this question, we used a carbon dioxide electrode, which was designed to measure the total amount of dissolved carbon dioxide in solution. The solution in the internal filling of the electrode is separated from the analytical sample by a CO_2_ permeable membrane. Typically, a carbon dioxide buffer is added to the analytical sample to lower the pH to 4.8-5.2 thus shifting the CO_2_/bicarbonate equilibrium 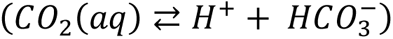 towards free CO_2_. Carbon dioxide diffuses through the membrane, and an equilibrium is reached between the concentration of CO_2_ in the analyte and the internal filling. The increase of CO_2_ in the internal filling causes an increase in hydrogen ion concentration, which is ultimately measured by the pH electrode. To measure the actual (equilibrium concentration) rather than the total concentration of carbon dioxide, we used this electrode in a sealed system and without the carbon dioxide buffer. To calibrate our measurement system, we compared the calculated concentrations of CO_2_ in sodium bicarbonate buffers with those obtained by direct measurements with the carbon dioxide electrode (Figure 5). The results showed a faithful relationship (two-tailed t-test, *P*=0.1037) between calculated and measured CO_2_ concentrations (Figure 5). We then used this system to measure the concentrations of dissolved CO_2_ in samples obtained by bubbling CO_2_ in perfusion Ringer buffer. The higher the concentration of CO_2_, the larger the currents recorded from CquiGR2/CquiGR3-coexpressing oocytes (Figure S3). Of note, currents recorded from sodium bicarbonate is the summation of carbon dioxide and Na^+^ responses. Although we adjusted the pH of the tested bubbled CO_2_ samples (Figure S3) to rule out the possible effect of hydrogen ion concentrations on oocyte responses, these recordings did not unambiguously determine whether the receptor system is responding to carbon dioxide per se or bicarbonate.

**Figure 5.**
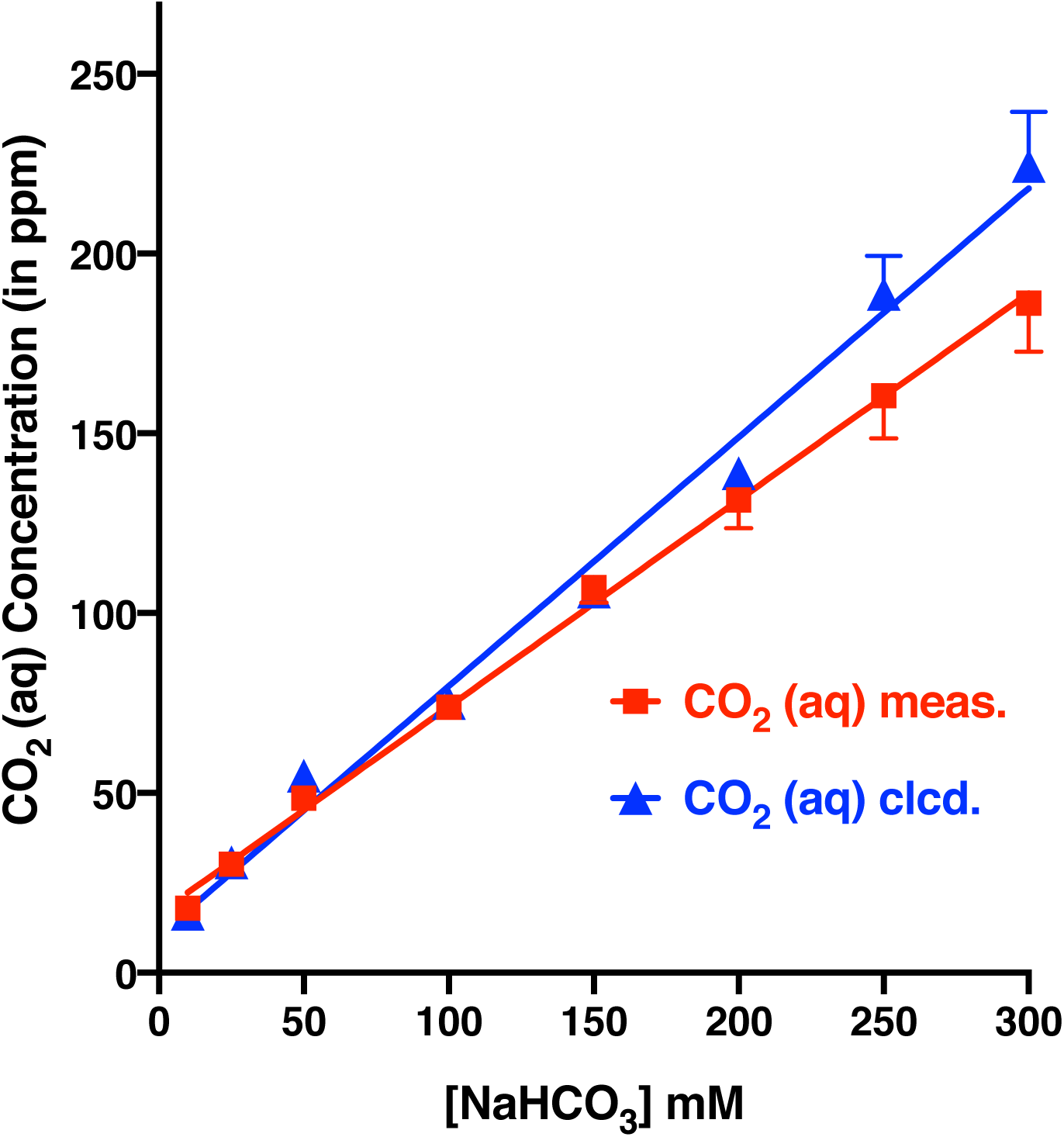
CO_2_ Concentrations in Solutions of Sodium Bicarbonate Prepared in Ringer Buffer. The calculated concentrations based on the final pH of each preparation (n *=* 4) are shown in red. The actual concentration of CO_2_ in each solution was measured with a CO_2_ Ion Selective Electrode. Data are presented as mean ± SEM.

The next experiment further examined the possible role of pH on recorded currents. We injected Ringer and 50 mM sodium chloride samples with their pH adjusted to that of the perfusion Ringer buffer, as well as to 7, 6.5, 6, and 5.5. To make certain that the CquiGR2/CquiGR3-coexpressing oocyte system was functional throughout the tests, we injected Ringer, sodium chloride, and sodium bicarbonate at the beginning and the end of each run (Figure S4). These experiments (n = 6) further demonstrated that hydrogen ion concentration at the pH range of 7.94 to 5.5 had little or no effect on sodium bicarbonate-elicited response (Figure S4).

Having developed tools to measure the amount of free carbon dioxide in solution, CO_2_ (aq), and demonstrated that pH change had minimal or no effect on oocyte responses, we next tested whether sifting the equilibrium of CO_2_/bicarbonate buffers would cause an increase or a decrease in the receptor response. We prepared in duplicates aliquots of 10 mM sodium bicarbonate buffers and placed them in sealed V-vials with Teflon liner, open-top caps. One of the experimenters took a sample from one of a pair of vials, applied to a CquiGR2/CquiGR3- coexpressing oocyte and recorded the elicited current. Meanwhile, the other experimenter injected an aliquot of 0.1 N HCl to the second pair of the same sample, vortexed, and provided the sample to the other experimenter to apply immediately to the oocyte preparation (Figure S5). The pairs of unadjusted and pH adjusted samples were kept sealed until their pH values were recorded. As expected, the initial pH was nearly the same (pH = 7.73) because they were aliquots from the same 10 mM sodium bicarbonate sample. The final pH of the tested samples was 7.39 ± 0.05.

Quantification of the recorded currents from paired samples showed a significant increase (two-tailed t-test, n *=* 12, p *=* 0.0005) in the lower pH samples (Figure 6A). Similar experiments were performed to measure the actual concentration of dissolved CO_2_ in paired samples with initial and lowered pH. The initial and final pH values of these samples (n *=* 4) were 7.73 and 7.32 ± 0.03, respectively. As expected, the quantification of CO_2_ (aq) showed an increase in dissolved CO_2_ at lower pH (Figure 6B). The measured CO_2_ concentrations in the samples without adjustment (pH_initial_) and with pH adjusted (pH_lowered_) were 17.90 ± 1.55 ppm and 32.40 ± 2.40 ppm, whereas the calculated CO_2_ concentrations were approximately 15.8 and 38.4 ppm, respectively. An increase in dissolved CO_2_ concentration by lowering pH is predicted by Le Chatelier’s Principle, given that an increase in H^+^ concentration shifts the equilibrium towards CO_2_ (aq). Because the increased oocyte response (Figure 6A) resulted from the increase of in situ CO_2_ concentration (Figure 6B), we concluded that CO_2_ per se, not bicarbonate, activates the *Cx. quinquefasciatus* gustatory receptor system when expressed in *Xenopus* oocytes. It has also been reported that the BAG neurons of the nematode *Caenorhabditis elegans* are activated by molecular CO_2_, although they can be activated by acid stimuli [22]. We found no evidence for the activation of the *Culex* mosquito GR system by hydrogen ions.

**Figure 6.**
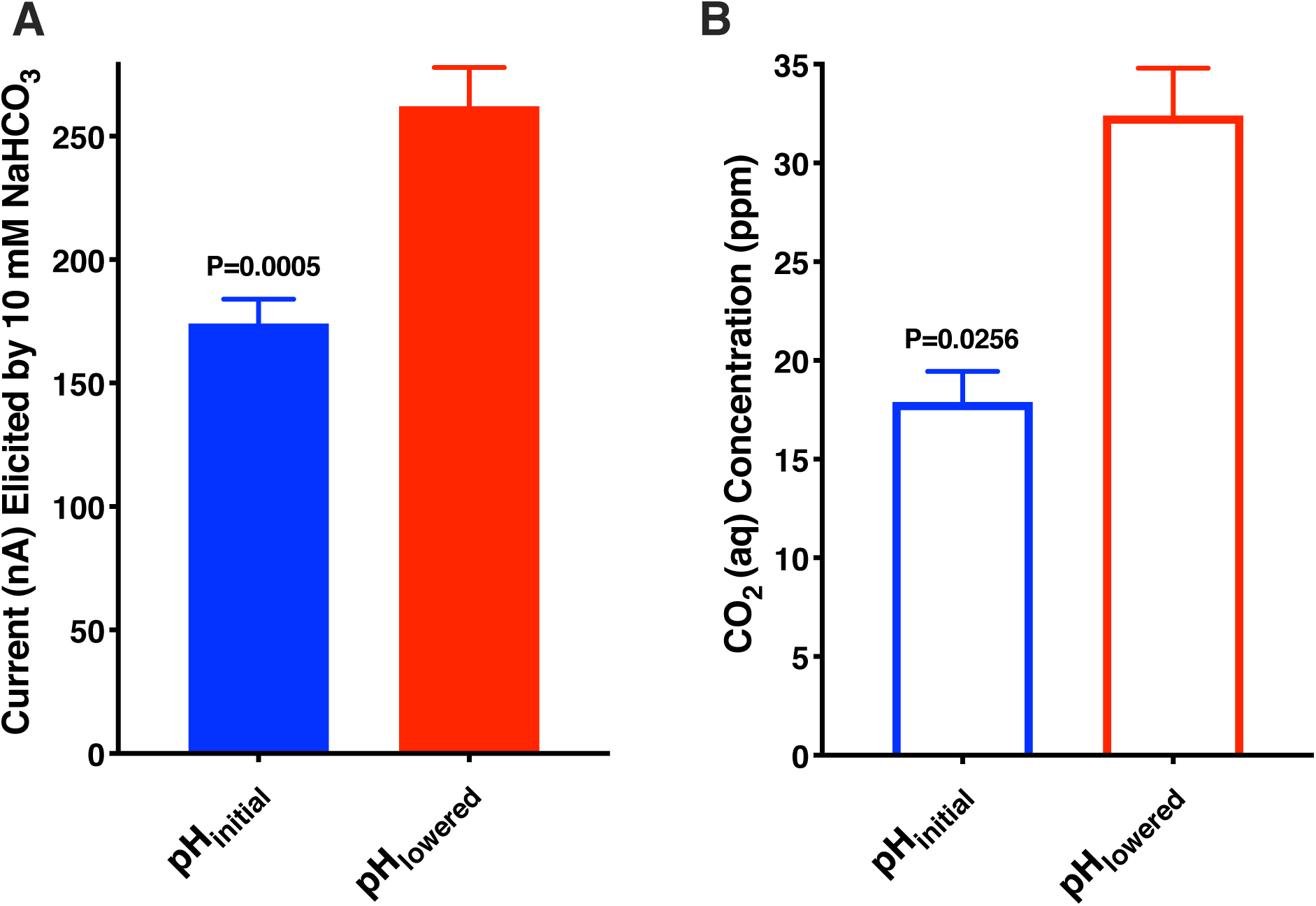
Effect of pH on the Oocyte Response and the Concentrations of CO_2_ in Solution. (A) Responses of CquiGR2+CquiGR3-expressing oocytes to 10 mM of sodium bicarbonate without acidification and after acidification (n = 12). The final pH (pH_lowered_) of the sodium bicarbonate buffer was 7.39 ± 0.05. (B) Quantification of CO_2_ in sodium bicarbonate samples (n *=* 4) without and after pH adjustment (acidification) as measured with a CO_2_ Ion Selective Electrode. After acidification (pH_lowered_), the pH of the sodium bicarbonate samples was 7.32 ± 0.03. Data are presented as mean ± SEM.

### CquiGR1 Orthologs in *An. gambiae* and *Ae. aegypti* are Putative Modulators

Given the unexpected results that CquiGR1 was not necessary for receptor function, but rather attenuated the responses of the CquiGR2/CquiGR3 receptor system to CO_2_, we asked whether this would be a common feature of mosquito biology. We prepared oocytes to coexpress the three GR subunits from the malaria mosquito, specifically AgamGR22, AgamGR23, and AgamGR24. Using the same batch of oocytes, we prepared a binary combination of these receptors devoid of AgamGR22, ie, AgamGR23/AgamGR24-expressing oocytes. Then we compared the responses of these oocytes to the same concentrations of sodium chloride and sodium bicarbonate. Specifically, we challenged each oocyte preparation of the same age and the same batch of oocytes with 50, 100, 200, and 300 mM solutions of NaCl and NaHCO_3_. AgamGR23/AgamGR24-coexpressing oocytes gave stronger, concentration-dependent responses to sodium bicarbonate (Figure 7A) than the oocytes coexpressing a tertiary receptor system, AgamGR22/AgamGR23/AgamGR24 (Figure 7B). These experiments were replicated with another batch of oocytes (Figure 7C,D). These results suggest that the CquiGR1 ortholog from the malaria mosquito, ie, AgamGR22, functions as a modulator.

**Figure 7.**
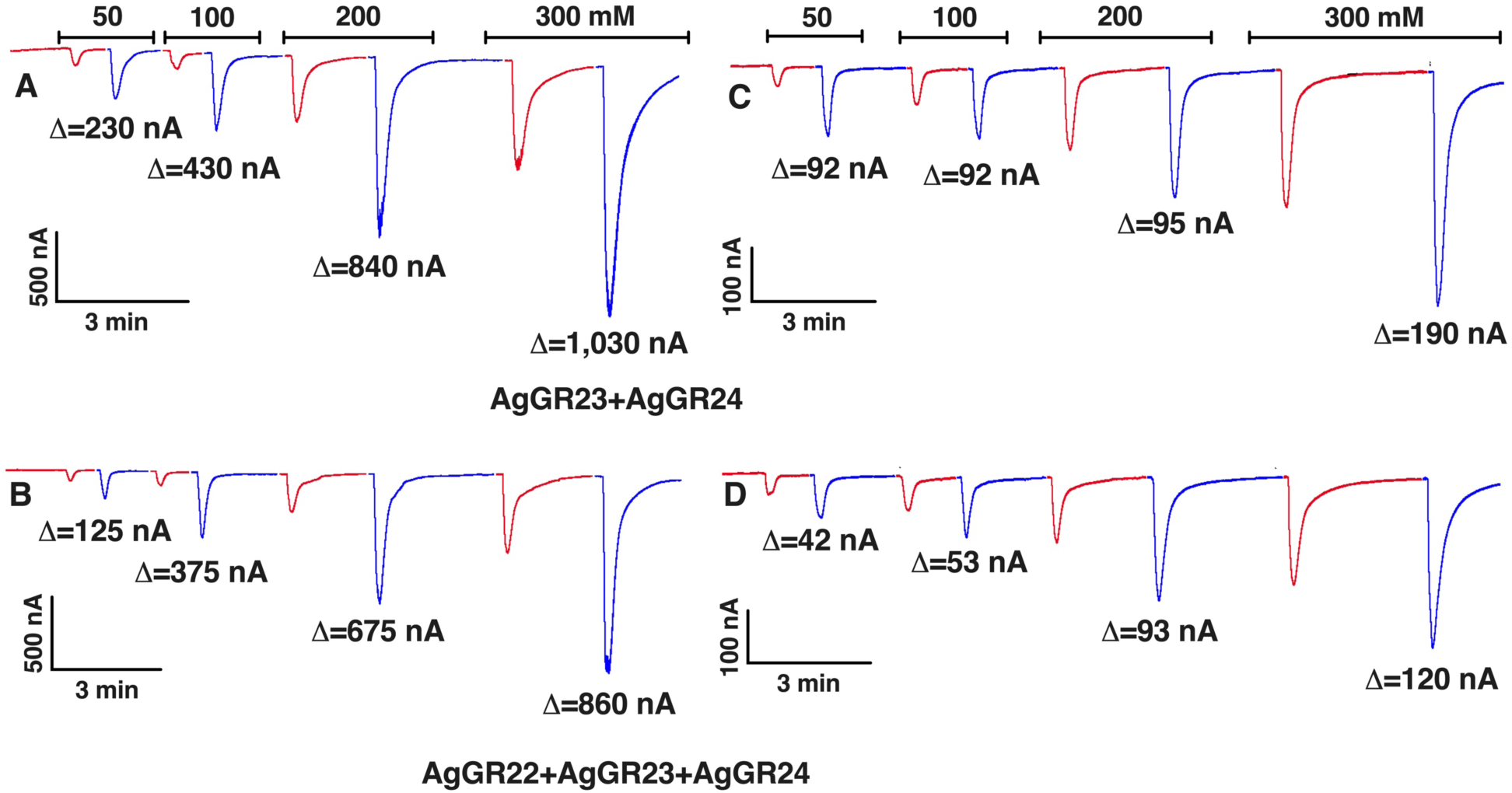
Responses of *An. gambiae* GRs-expressing Oocytes to Sodium Chloride and Sodium Bicarbonate. (A) Trace obtained with an oocyte coexpressing AgamGR23 and AgamGR24. (B) Responses of another oocyte (from the same batch) coexpressing the three GRs from *An. gambiae*, GR22, GR23, and GR24. (C) Replication of experiment in (A) but using a different batch of oocytes. (D) Replication of the experiment in (B) with the same batch of oocytes used in (C). Δ values represent responses to sodium bicarbonate subtracted from responses to sodium chloride at the same concentration. In a third replication with a different batch of oocytes the Δ values recorded from a GR23+GR24-coexpressing oocyte stimulated with 50, 100, 200, and 300 mM were 120, 225, 250, and 780 nA, whereas the respective values recorded from an oocyte from the same batch and coexpressing GR22+GR23+GR24 were 23, 89, 193, and 270 nA.

Next, we prepared oocytes coexpressing the equivalent binary and tertiary receptor systems with GRs from the yellow fever mosquito. Specifically, we compared the responses of AaegGR2/AaegGR3-coexpressing oocytes to oocytes of the same batch and the same age that coexpressed AaegGR1, AaegGR2, and AaegGR3. Again, the receptor system devoid of AaegGR1 generated stronger, dose-dependent responses to bicarbonate (Figure S6A) than the tertiary system (Figure S6B) thus implicating AaegGR1 in a modulatory role.

## CONCLUSIONS

Our research plans to address a long-standing question regarding CO_2_ reception led to unexpected results. Because it has been well-established in the literature that the reception of CO_2_ in the vinegar fly antennae is mediated by DmelGR21a and DmelGR63a [12–14], it has been assumed that CO_2_ is detected by GR1+GR3 in mosquitoes. This notion has been supported by RNAi-based experimental data suggesting that GR2 was not involved in CO_2_ reception in *Ae. aegypti* (and *Cx. quinquefasciatus*) [21]. On the other hand, AgamGR23, the CquiGR2 ortholog from the malaria mosquito *An. gambiae*, was implicated in CO_2_ response [20], but the role of AgamGR22 is yet to be unraveled. Our findings suggest that the three subunits might be involved, with GR1 (GR22 in the case of *An. gambiae*) being a putative modulator. With a functional carbon dioxide-detecting receptor system in an aqueous environment, we applied a fundamental principle in chemistry to interrogate which of two species in equilibrium (CO_2_ or HCO_3_^-^) might activate the receptors. We compared receptor responses as well as CO_2_ concentrations in paired samples without acidification and after acidification and showed that an increase in the concentrations of dissolved CO_2_ was associated with an increase in receptor response. We then concluded that CO_2_ per se, neither HCO_3_^-^ nor protons, activates the carbon dioxide-detecting system in mosquitoes.

## STAR METHODS

All methods can be found in the accompanying Transparent Methods supplemental file.

## SUPPLEMENTAL INFORMATION

Supplemental information includes Transparent Methods and six supplementary figures that can be found with this article online at https……

## ACKNOWLEDGMENTS

We thank Dr. Su Liu for comments on a draft version of the manuscript. X.W. was supported by the Chinese Scholarship Council. Research reported in this publication was supported by the National Institute of Allergy and Infectious Diseases (NIAID) of the National Institutes of Health under award number R01AI095514. The content is solely the responsibility of the authors and does not necessarily represent the official views of the NIH.

## AUTHOR CONTRIBUTIONS

P.X., X.W., and W.S.L. performed research; W.S.L. designed research; P.X. and WS.L. analyzed the data; W.S.L. wrote the paper; all authors read and approved the final manuscript.

## DECLARATION OF INTEREST

The authors declare no conflict of interest.

Received: …..

---

**Figure S1.**
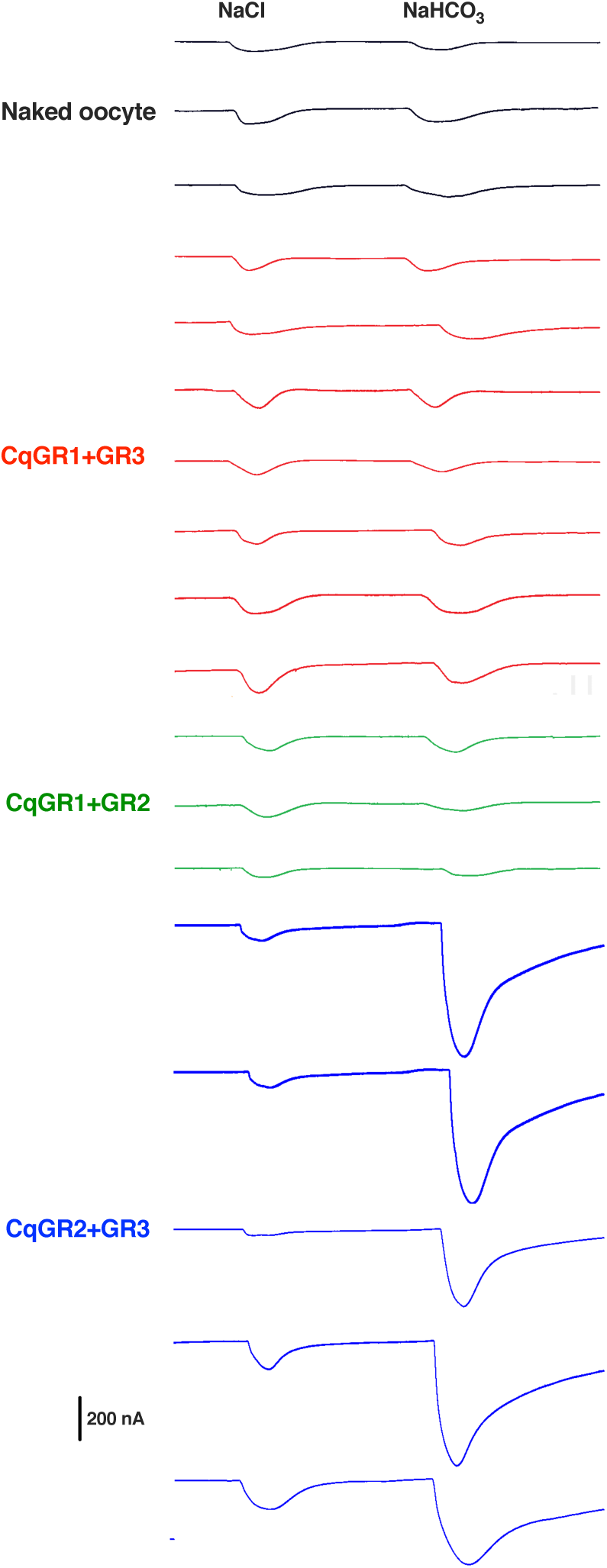
NaCl and NaHCO_3_ Responses Recorded from Oocytes only and Oocytes Coexpressing combination of *Culex* GRs. From top to bottom: naked oocytes (black), oocytes coexpressing GR1+GR3 (red), GR1/GR2- coexpressing oocytes (green), and oocytes coexpressing GR2 and GR3 (blue). NaCl and NaHCO_3_ were applied at 200 mM.

**Figure S2.**
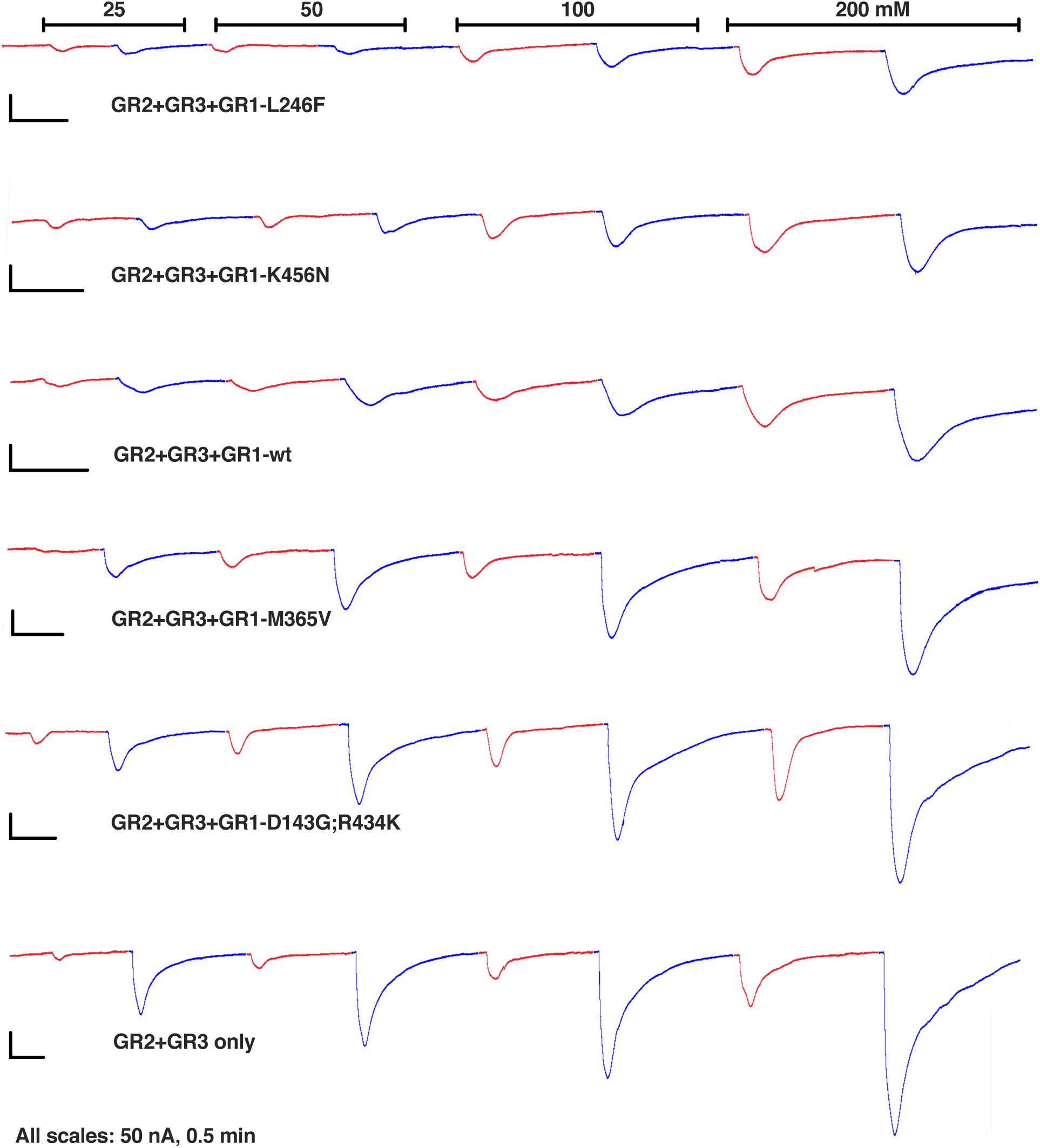
Effect of CquiGR1 Variants on the Modulation of CO_2_ Response. Traces obtained with the same batch of oocytes injected either with CquiGR2+CquiGR3 or a tertiary combination of CquiGR2+CquiGR3 plus one of the CquiGR1 variants. For clarity, responses to sodium chloride and sodium bicarbonate are colored red and blue, respectively.

**Figure S3.**
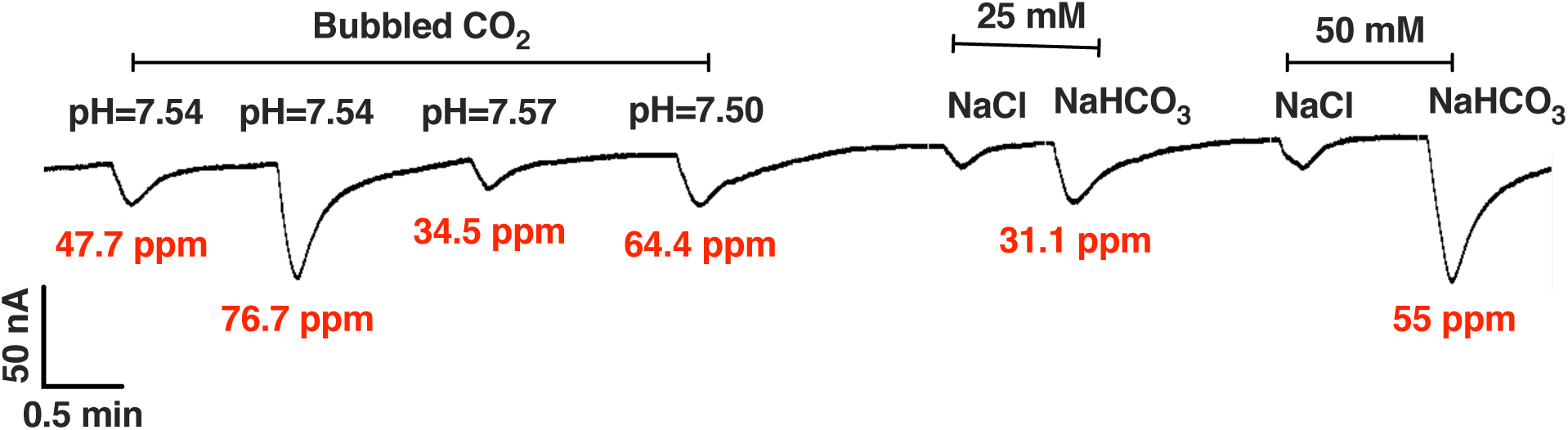
Responses Obtained with a CquiGR2+CquiGR3-expressing Oocyte when Stimulated with Carbon Dioxide. Samples were prepared by bubbling CO_2_ in Ringer buffer and adjusting the final pH or by dissolving either sodium chloride or sodium bicarbonate in Ringer buffer. The concentrations of CO_2_ (in samples prepared by bubbling CO_2_ or by dissolving NaHCO_3_) were measured using a CO_2_ Ion Selective Electrode. Of note, currents elicited by sodium carbonate are larger than expected for the amount of CO_2_ (aq), because they include currents elicited by sodium.

**Figure S4.**
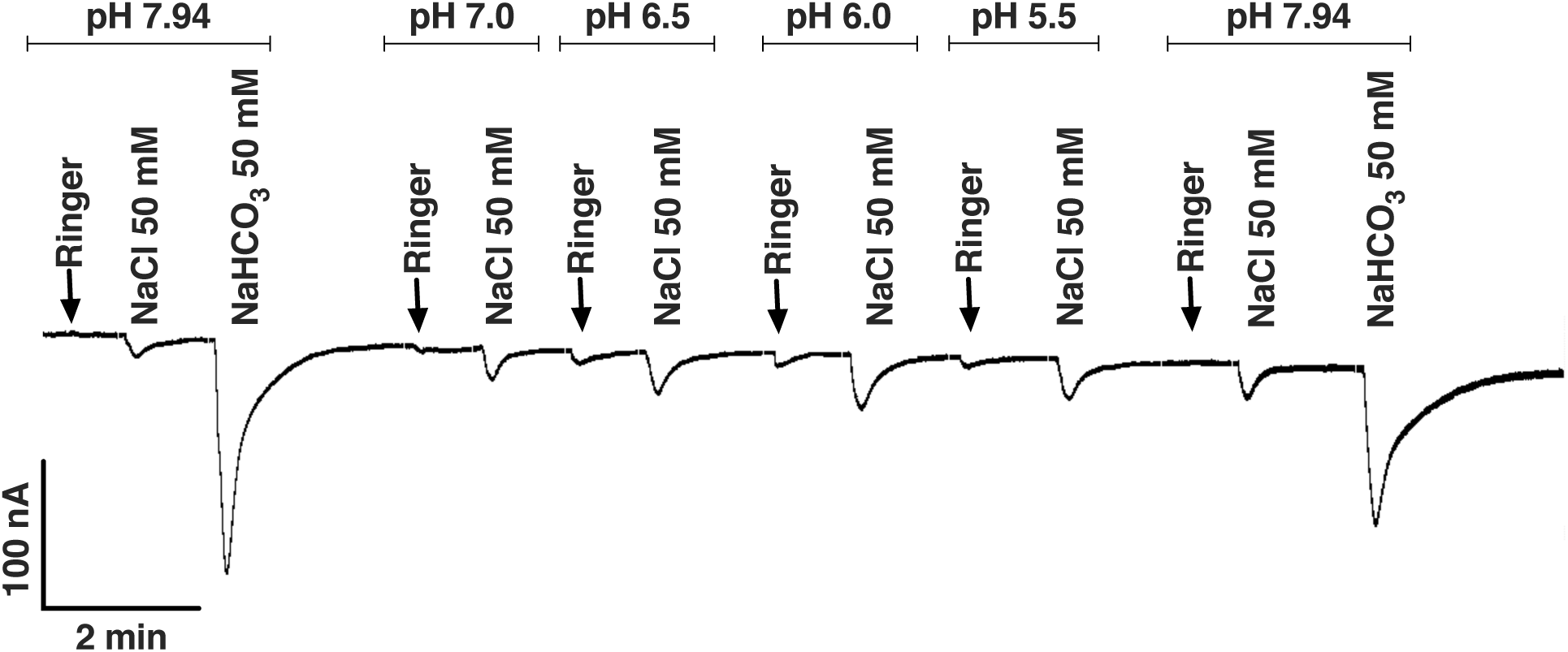
A Continuous Trace Obtained with an Oocyte Coexpressing CquiGR2+CquiGR3. In this experiment, a freshly prepared perfusion Ringer buffer (pH 7.94) was used. Ringer buffer and 50 mM samples of sodium chloride or sodium bicarbonate at the same pH were applied at the beginning and the end of the experiment. The pH values of Ringer buffers and 50 mM NaCl solutions were adjusted to 7, 6.5, 6, and 5.5 before injection. These experiments were repeated six times.

**Figure S5.**
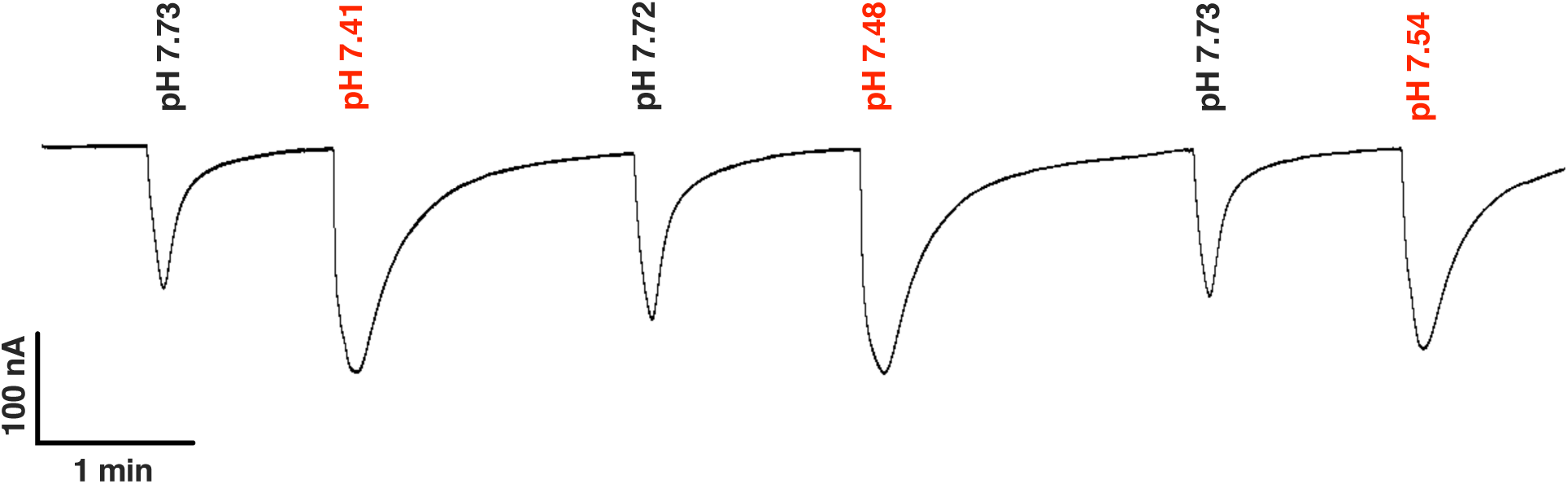
Trace Obtained from an oocyte coexpressing CquiGR2 and CquiGR3. The pH of injected samples (10 mM NaHCO_3_) was measured before and after acidification.

**Figure S6.**
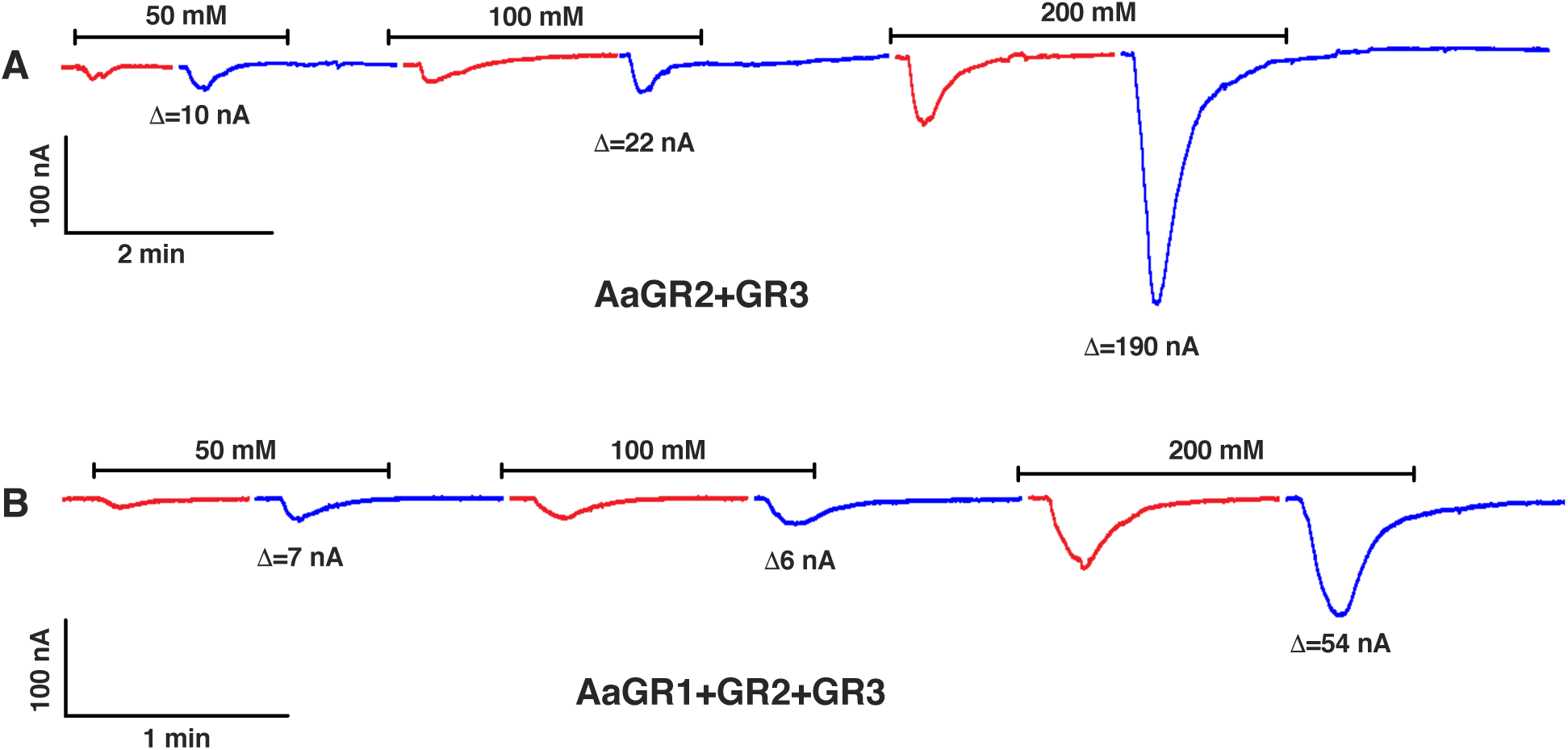
Responses of *Ae. aegypti* GRs Co-expressed in *Xenopus* Oocytes and Challenged by Sodium Chloride and Sodium Bicarbonate. (A) Traces obtained with an AaegGR2/AaegGR3-coexpressing oocyte. (B) Responses recorded from another oocyte of the same age and the same batch, but coexpressing the subunits AaegGR1, AaegGR2, and AaegGR3. Δ values represent responses to sodium bicarbonate subtracted from responses to sodium chloride at the same concentration.

## TRANSPARENT METHODS

### RNA extraction, DNA synthesis, and cloning

Three hundred pairs of palps from 4- to 6-d-old female *Culex* mosquitoes were dissected under a stereomicroscope (Zeiss, Stemi DR 1663) and collected in 75% (vol/vol) ethanol diluted in diethylpyrocarbonate (DEPC)-treated water on ice. The samples were centrifuged, and ethanol was removed before total RNA was extracted using TRIzol reagent (Invitrogen). cDNA was synthesized from 1 μg of total RNA using GoScript™ Reverse Transcription kit according to the manufacturer’s instructions (Promega). For *An. gambiae* and *Ae. aegypti*, gene sequences of AgamGR22, 23, 24 and AaegGR1, 2, 3 from Genbank were synthesized by GenScript USA Inc. (Piscataway, NJ). Full-length gene-specific cloning primers were designed based on the sequences from Genbank and added 15-bp In-Fusion cloning adapters.

CqGR1-Fw <u>GATCAATTCCCCGGG</u>accATGATTCACAGTCAGATGGAGGACG

CqGR1-Rv <u>CAAGCTTGCTCTAGA</u>CTATTGTGGGTTCGTTTTGGCCG

CqGR2-Fw <u>GATCAATTCCCCGGG</u>accATGGTCATCAAGGACAGTGACTTTGAC

CqGR2-Rv <u>CAAGCTTGCTCTAGA</u>TTAGTGAGCGTGAGCTTTCTGTAAATTCTC

CqGR3-Fw <u>GATCAATTCCCCGGG</u>accATGAGCATATTTCCGGATACTCTGCG

CqGR3-Rv <u>CAAGCTTGCTCTAGA</u>TCAATGCGCTGCCGTCG

AgGR22-Fw <u>GATCAATTCCCCGGG</u>accATGATTCACACACAGATGGAAG

AgGR22-Rv <u>CAAGCTTGCTCTAGA</u>TTAGGTGTTCACTTTGTCTGC

AgGR23-Fw <u>GATCAATTCCCCGGG</u>accATGGTTATCAAGGAAAGTGAGTTC

AgGR23-Rv <u>CAAGCTTGCTCTAGA</u>TTACTGTTTGTGTAGCAGCTTAACA

AgGR24-Fw <u>GATCAATTCCCCGGG</u>accATGAGTCTCTACTTCAACGCGG

AgGR24-Rv <u>CAAGCTTGCTCTAGA</u>CTAAGAATGAGACGAATTACTGTGC

AaGR1-Fw <u>GATCAATTCCCCGGG</u>accATGATTCACAGCCAGATGGAAG

AaGR1-Rv <u>CAAGCTTGCTCTAGA</u>CTAGTTCTCCTTCAGCTTAGTTAGA

AaGR2-Fw <u>GATCAATTCCCCGGG</u>accATGGTCATCAAAGACAGTGAGT

AaGR2-Rv <u>CAAGCTTGCTCTAGA</u>CTATCCCTTATGACTGTGCTTGATT

AaGR3-Fw <u>GATCAATTCCCCGGG</u>accATGAATCTCAACCAAGACCCCA

AaGR3-Rv <u>CAAGCTTGCTCTAGA</u>CTACTCGCGATATGAACCCGTCATA

Adapter sequences are underlined, and lower case acc is Kozak consensus. PCR products were purified by Monarch^®^ DNA Gel Extraction Kit (NEBioLab) and directly inserted to linearized destination vector-pGEMHE by In-Fusion reaction (Clontech). The colonies were picked, and vectors were extracted using Monarch^®^ Plasmid Miniprep Kit. Inserts were submitted to DNA Sequencing Facility, UC Berkeley for verification of sequences.

### Oocytes preparations and two-electrode voltage clamp recordings

The vectors carrying GR genes were linearized with restriction endoneuclease NheI, except for AgamGR22, which was cut by SphI. The linearized vectors with gene inserts were used as templates to transcribe capped cRNAs with poly(A) using an mMESSAGE mMACHINE T7 kit (Ambion) following the manufacturer’s protocol. The cRNAs were dissolved in RNase-free water and adjusted at a concentration of 200 μg/mL by UV spectrophotometry (NanoDrop™ Lite Spectrophotometer, ThermoFisher). 9.2 nl of each cRNA samples were microinjected into stage V or VI *Xenopus* oocytes (purchased from EcoCyte Bioscience, Austin, TX). Then injected oocytes were incubated at 18 °C for 3–7 days in modified Barth’s solution [in mM: 88 NaCl, 1 KCl, 2.4 NaHCO_3_, 0.82 MgSO_4_, 0.33 Ca(NO_3_)_2_, 0.41 CaCl_2_, 10 HEPES, pH 7.4] supplemented with 10 μg/mL of gentamycin, 10 μg/mL of streptomycin, and 1.8 mM sodium pyruvate. For two-electrode voltage clamp (TEVC) recordings, as previously done [1], oocytes were placed in a perfusion chamber and challenged with test compounds. Compound-induced currents were amplified with an OC-725C amplifier (Warner Instruments, Hamden, CT), the voltage held at −80 mV, low-pass filtered at 50 Hz and digitized at 1 kHz. Data acquisition and analysis were carried out with Digidata 1440A and pClamp10 software (Molecular Devices, LLC, Sunnyvale, CA).

### Sample preparations and CO_2_ measurement

Sodium chloride (Fisher Scientific, >99%) and sodium bicarbonate (Sigma-Aldrich, 99.7%) were used to prepare fresh samples by dissolving the appropriate amounts of these salts in perfusion Ringer buffer (NaCl 96 mM, KCl 2 mM, CaCl_2_ 1.8 mM, MgCl_2_ 1 mM, HEPES 5 mM, pH 7.6) to make 0.5 M solutions. Then, the desired concentrations were prepared by diluting with perfusion Ringer buffer. CO_2_ samples were prepared by bubbling MediPure™ CO_2_ (Praxair, Danbury, CT) directly into perfusion buffer for 5 s, and subsequently adjusting the pH.

The concentration of dissolved CO_2_, ie, CO_2_ (aq), was calculated by equation I:

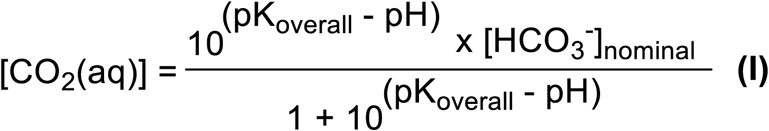

K_overall_, which is sometimes referred to as K_a_, is a constant, which incorporates the constant of hydrolysis of CO_2_ and the first dissociation constant of carbonic acid, K_a1_, ie, K_overall_ = K_h_ x K_a1_. pKa is 6.3 (https://pubchem.ncbi.nlm.nih.gov/compound/Sodium-bicarbonate#section=pKa). The molar concentration of dissolved CO_2_ is, therefore, related to the pH of the solution and the nominal concentration of sodium bicarbonate. Thus, the concentration of dissolved CO_2_ in a 10 mM NaHCO_3_ solution at pH 7.73 is approximately 0.358 mM or 15.76 ppm. At pH 7.32, the calculated concentration of dissolved CO2 is 0.872 mM or 38.36 ppm.

pH and carbon dioxide were measured with an Orion Star™ A214 pH/ISE Benchtop Meter (ThermoFisher Scientific, Waltham, MA 02451). A carbon dioxide electrode was used for CO_2_ measurements. The internal filling solution of this electrode is separated from the sample solution with a gas-permeable membrane. CO_2_ dissolved in a sample solution diffuses through this membrane until an equilibrium between CO_2_ in the electrode filling and the sample solution is reached. The influx of CO_2_ causes a change in hydrogen ion concentration in the electrode filling solution, which is measured by a sensing element behind the membrane. The system was calibrated before measurements with freshly prepared 22, 55 and 110 ppm solutions. Standard solutions were sealed with parafilm before measurements. To minimize losses of carbon dioxide during measurements, we used 50 ml Falcon tubes to house sample solutions. A hole was opened in the tube cap, and the electrode was inserted and kept in place with two O-ring seals below and two O-ring seals on the top of the cap. Then the cap was sealed with Teflon tapes and finally covered with electrical tape. Fifty milliliter Falcon tubes housing test samples were attached to the electrode and fastened tightly. The concentrations of dissolved CO_2_ in bicarbonate solutions were measured by preparing 10, 25, 50, 100, 150, 200, 250, and 300 mM solutions in perfusion Ringer buffer, with four replicates for each sample. To compare the concentrations of dissolved CO_2_ without adjustment and after acidification, 7 ml aliquots of 10 mM sodium bicarbonate buffers were prepared in pairs. One of the two aliquots in a pair was used to determine CO_2_ concentration without pH change. To the other pair, 150 µl of 0.1 N HCl was added and immediately used for CO_2_ measurement, and then the pH was recorded. To compare oocyte response, samples were similarly prepared by using smaller aliquots (700 µl) in V-vials. In this case, 0.1 N HCl was injected through the Teflon liner of the open-top caps. After that, an aliquot was collected to be applied to the *Xenopus* oocyte recording system, and the remainder was used for pH measurement.

### Statistical analysis and graphical preparations

Prism 8.2.0 from GraphPad Software (La Joya, CA) was used for both statistical analysis and graphical preparations. A dataset that passed the Shapiro-Wilk normality test was analyzed by t-test; otherwise, data were analyzed by Wilcoxon matched-pairs signed-rank test. All data are presented as mean ± SEM.

